# *Fgf8* regulates first pharyngeal arch segmentation through pouch-cleft interactions

**DOI:** 10.1101/2023.03.15.532781

**Authors:** Nathaniel Zbasnik, Jennifer L. Fish

## Abstract

The pharyngeal arches are transient developmental structures that, in vertebrates, give rise to tissues of the head and neck. A critical process underlying the specification of distinct arch derivatives is segmentation of the arches along the anterior-posterior axis. Out-pocketing of the pharyngeal endoderm between the arches is a key mediator of this process, and although it is essential, mechanisms regulating out-pocketing vary between pouches and between taxa. Here, we focus on the patterning and morphogenesis of epithelia associated with the first pharyngeal arch, the first pharyngeal pouch (pp1) and the first pharyngeal cleft (pc1), and the role of *Fgf8* dosage in these processes. We find that severe reductions of *Fgf8* levels disrupt both pp1 and pc1 development. Notably, out-pocketing of pp1 is largely robust to *Fgf8* reductions, however, pp1 extension along the proximal-distal axis fails when *Fgf8* is low. Our data indicate that extension of pp1 requires physical interaction with pc1, and that multiple aspects of pc1 morphogenesis require *Fgf8*. In particular, *Fgf8* is required for specification of regional identity in both pp1 and pc1, for localized changes in cell polarity, and for elongation and extension of both pp1 and pc1. Overall, our data indicate a critical role for the lateral surface ectoderm in segmentation of the first pharyngeal arch that has previously been under-appreciated.

## INTRODUCTION

The pharyngeal arches (PAs) are transient developmental structures that, in vertebrates, give rise to tissues of the head and neck. The PAs are formed by mesenchymal populations, the neural crest (NC) and mesoderm, that migrate in between epithelial layers, the surface ectoderm and foregut endoderm (Depew, 2002). The mesenchymal cells of each arch give rise to specific musculoskeletal derivatives. For example, the jaw derives from the first pharyngeal arch (PA1), while the second arch, in mammals, forms the stapes and contributes to the hyoid bone. A critical process underlying the specification of these different skeletal elements is segmentation of the arches along the anterior-posterior axis, which ensures separation of mesenchymal populations and allows for their differential patterning (Graham and Smith, 2001). Arch segmentation is mediated by the pharyngeal epithelia. Pharyngeal pouches form via localized out-pocketing of the foregut endoderm that grows laterally to contact the ectoderm, which invaginates to form the pharyngeal clefts (Graham, 2001, 2003; Graham *et al*., 2005). Mutations or experimental alterations resulting in the failure to generate pharyngeal pouches also cause failure of pharyngeal arch formation (Couly *et al*., 2002; Edlund *et al*., 2014; Piotrowski and Nusslein-Volhard, 2000).

In addition to arch segmentation, pharyngeal epithelia provide essential signals regulating the patterning and proliferation of arch mesenchyme (Brito *et al*., 2006; Couly *et al*., 2002; Edlund *et al*., 2014; Hasten and Morrow, 2019; Haworth *et al*., 2007; Trumpp *et al*., 1999). Subsequent to these key patterning roles, the pouches and clefts also give rise to essential tissues of the head (Grevellec and Tucker, 2010). Thus, the pharyngeal epithelia have at least 3 critical roles: 1) segmenting the arches, 2) providing signals that support patterning and proliferation of arch mesenchyme, and 3) differentiating into tissue derivatives. Given that differential patterning of each arch along the anterior-posterior axis is required for the formation of arch-specific derivatives (e,g., Gendron-Maguire *et al*., 1993; Rijli *et al*., 1993; Trainor and Krumlauf, 2001), and tissue derivatives of the pouches and clefts differ among vertebrate taxa (Grevellec and Tucker, 2010; Poopalasundaram *et al*., 2019), it is not surprising that arch segmentation varies between arches along the anterior-posterior axis and between the same arch in different taxa. These differences have been characterized in terms of ectodermal–endodermal tissue relationships (Shone and Graham, 2014) and molecular mechanisms of out-growth (Choe *et al*., 2013; Crump *et al*., 2004; Hasten and Morrow, 2019; Okubo *et al*., 2011; Piotrowski *et al*., 2003; Quinlan *et al*., 2002).

Our work investigates development of the lower jaw and how alterations to jaw development contribute to human disease. Therefore, we specifically focus on the patterning and morphogenesis of PA1 and its associated epithelia, the first pharyngeal pouch (pp1) and the first pharyngeal cleft (pc1). We previously described how *Fgf8* has a dosage effect on jaw size. Specifically, reductions in *Fgf8* expression below about 40% of WT expression levels result in truncated and dysmorphic lower jaws (Green *et al*., 2017; Zbasnik *et al*., 2022). Mild mutants (*Fgf8^Neo/Neo^*; expressing 35% of *Fgf8* relative to WT) exhibit minor truncations of the proximal mandible whereas more severe mutants (*Fgf8^Δ/Neo^*; expressing 20% of *Fgf8* relative to WT) exhibit unilateral fusion of the upper and lower jaws (Zbasnik *et al*., 2022). Importantly, these defects are associated with malformations of PA1 epithelia which fail to separate the first and second arches. As a consequence, the expression patterns of key patterning genes are altered. Based on these data, we hypothesized that *Fgf8*-mediated morphogenesis of pharyngeal epithelia is critical to arch segmentation and PA1 patterning.

Defects in pp1 morphogenesis are associated with several craniofacial disease syndromes including DiGeorge (22q11.2 deletion) Syndrome (Arnold *et al*., 2006; Frank *et al*., 2002; Jerome and Papaioannou, 2001; Moon *et al*., 2006; Piotrowski *et al*., 2003; Zhang *et al*., 2005). Although human FGF8 does not localize to 22q11, *Fgf8* deficiency in mice generates many features of 22q11.2 deletion syndromes (Frank *et al*., 2002). Fgf8 is an important developmental signaling factor that genetically interacts with *Tbx1* and *Crkl*, two genes that lie within 22q11.2 (Moon *et al*., 2006; Vitelli *et al*., 2002a; Vitelli *et al*., 2002b). *Fgf8* is expressed in both pp1 and pc1 where it overlaps with *Tbx1* and *Foxi3* expression (Hasten and Morrow, 2019). Both *Tbx1* and *Foxi3* have been reported to be upstream of *Fgf8* (Edlund *et al*., 2014; Hasten and Morrow, 2019; Vitelli *et al*., 2002b). In *Foxi3^-/-^* embryos, *Fgf8* expression is greatly reduced in the ectoderm and endoderm, pc1 and pp1 do not form, and PA1 and PA2 fail to separate (Edlund *et al*., 2014). Interestingly, tissue-specific deletion of *Fgf8* in pp1 does not affect its morphogenesis or craniofacial development (Jackson *et al*., 2014). Extensive evidence supports a critical role for *Fgf8* expression, particularly from the oral ectoderm, in inducing PA1 mesenchymal gene expression regulating jaw patterning (e.g., Ferguson *et al*., 2000; Fish *et al*., 2011; Griffin *et al*., 2013; Trumpp *et al*., 1999; Tucker *et al*., 1998). The role of *Fgf8* expression in epithelial patterning and morphogenesis is less understood. We therefore examined pharyngeal epithelial morphogenesis, focusing on pp1 and pc1, in embryos of varying *Fgf8* dosage.

The pharyngeal pouches develop serially from the primitive gut tube along the rostral-caudal axis (Graham *et al*., 2005; Veitch *et al*., 1999). Pouch morphogenesis relies on two separate morphogenetic processes, lateral out-pocketing and proximal-distal extension (Graham and Smith, 2001; Shone and Graham, 2014). Lateral out-pocketing is the process where the endoderm migrates laterally to contact the overlying ectoderm. Proximal-distal extension is the directional expansion of the pouch towards the midline of the embryo. Using an allelic series of *Fgf8* mice (Meyers *et al*., 1998), we characterized these processes in pp1 at varying *Fgf8* doses. We found that interaction between pp1 and pc1 is required for proximal-distal extension of pp1 and separation of PA1 and PA2. *Fgf8* is required for the interaction between pp1 and pc1 as this epithelial interface is disrupted in *Fgf8^Δ/Neo^* embryos and the arches fail to separate distally. Our data highlights the importance of the first ectodermal cleft in jaw patterning and suggests it may be a target of alteration in both disease and evolution.

## METHODS

### Experimental animals

The *Fgf8* allelic series utilizes 3 adult genotypes that can be crossed to generate five different embryonic genotypes (Meyers *et al*., 1998). Quantification of Fgf8 mRNA levels in E10.5 heads indicates that mice heterozygous for the Neo allele (*Fgf8^Neo/+^*) express 90% of *Fgf8* (wildtype; WT) levels, mice heterozygous for the Delta allele (*Fgf8^Δ/+^*) express 60% of WT levels, mice homozygous for the Neo allele (*Fgf8^Neo/Neo^*) express 35% of WT levels, and compound mutants (*Fgf8^Δ/Neo^*) express 20% of WT levels (Green *et al*., 2017). These mice and embryos were genotyped as previously reported (Green *et al*., 2017; Meyers *et al*., 1998; Zbasnik *et al*., 2022). Embryos were collected and staged based on the number of days after the observation of a postcoital plug at E0.5. Mouse experiments were approved by the University of Massachusetts Lowell Institutional Animal Care and Use Committees. Embryos were dissected on ice and fixed in 4% paraformaldehyde.

### *in situ* hybridization

Probes for *in situ* hybridization were generated from RNA isolated from E9.5 and E10.5 embryos. cDNA was produced using the Maxima first strand synthesis kit (ThermoFisher; K1641). The cDNA was then used as a PCR template to amplify the gene of interest (GOI). Select PCR primers had linkers (Fw:5’-ggccgcgg-3’; Rv:5’ - cccggggc-3’) to allow for a nested PCR TOPO cloning. PCR purification of these templates used a gene specific forward primer (GSFP) and the T7 Universal primer to amplify the initial GOI template. Primers lacking linkers were TOPO cloned into TOP10 competent E. coli cells using a PCR4 topo vector (ThermoFisher). Colonies were screened to ensure correct band length. Once purified, samples were sequenced to ensure the correct GOI was amplified and direction of the insertion. A second PCR was used to amplify the GOI using M13 primers after plasmid verification. Second PCR product lengths were verified on a 1% agarose gel. Products from the secondary PCR were used as the template for the anti-sense mRNA probe. Probes were made by using a dig RNA labeling mix (Sigma-Aldrich; #11277073910). Fluorescent probes were created with A555 fluorescent tyramine following publicly available protocols (https://sites.google.com/site/helobdellaprotocols/histology/tsa). Fluorescent samples were counter-stained using Hoechst’s O.N. (10 ug/mL) and were imaged using a 40x oil immersion objective lens on a Leica sp8 confocal microscope.

### Immunofluorescence

Whole embryos were fixed in 4%PFA/PBS, permeabilized with 0.1% Triton-X-100/PBS and then blocked in 5% FBS supplemented with 0.1% BSA for 1 hour. Primary antibodies used were: AP-2α (Santa Cruz sc-12726, 1:200 mouse), E-Cadherin (BD Transduction Laboratories 610181, 1:150 mouse), Laminin (Sigma-Aldrich L9393, 1:60 rabbit), and Sox2 (Millipore-Sigma AB5603-25UG, 1:1000 rabbit). The mitotic marker Phospho-Histone H3/ Ser10 (Cell Signaling 9701, 1:100 rabbit) was used to label proliferating cells. Secondary antibodies used against mouse were: Alexa488 (Abcam ab150113), Alexa647 (Invitrogen A-21240), Cy3 (Life Technologies A-10524) and against rabbit were: Alexa488 (Molecular Probes A11034), Alexa647 (Molecular Probes A21245). In conjunction with these antibodies, dyes were used to stain nuclei (Hoechst 1:1000), lysosomes to track cellular death (lysotracker Red) and F-actin cables (Phalloidin 1:1000). Embryos were imaged on a Nikon AR-1, a Leica Sp8, or a Zeiss Axiovert 200M microscope. Images were processed using the Fiji distribution of ImageJ and Adobe Photoshop CC 2018.

### Pouch shape analysis

Whole mount fluorescent *in situ* hybridization was performed for *Pax1*. Confocal images were collected at the point which the pouch opens and maintains contact with the cleft. Each side of the embryo was imaged through the total length of both the cleft and pouch until the first and second pouch connected. All left facing images were flipped along their vertical axis before landmarks were placed. 8 landmarks with 8 curves (semi landmark locations) were chosen to be digitized of each image for morphometric analyses. Stereomorph (version 1.6.3) a package within R (version 1.2.1335) was used to place these landmarks on each image. Once completed, the geomorphic package (version 3.3.2) was used to analyze the morphology of the first pouch.

A generalized procrustes analysis of the coordinates was performed using the gpagen function (Gower, 1975; Rohlf, 1990). The newly rotated and scaled shape data were then analyzed via procrustes ANOVA (procD.lm function). Additionally, a null model was made to verify if our chosen explanatory variables were contributing to shape changes. The null model was a fitted procrustes ANOVA of the shape coordinates with only the log transformed centroid size as a predictor variable. The full model ran against this null model included the log transformed centroid size, genotype, side, and somite number as predictor variables with all their interaction terms. An ANOVA was performed between the full and null model to verify if these two models were different. Finally, a principal component analysis (PCA) was performed on our coordinate data to generate a representative graph of pp1 shape change between *Fgf8* dosage levels. The gm.prcomp function was used to generate the graphs. Only younger (<23 somite pairs) and older (>28 somite pairs) embryos were used for the PCA analysis to maximize pp1 shape differences due to developmental age while ensuring we had enough samples in each group.

### Quantification of out-pocketing depth

Imaging parameters were the same as the shape analysis, with a z-step set to 2.5um. Each pouch was imaged from the point where the pp1 connects to pp2 medially through its full lateral extension. Depth was quantified by multiplying the number of frames by 2.5um. To test for significance, a general linear model was performed. Pouch depth was measured against the following fixed predictor variables: somite pairs (age), left or right side, and genotype. Somite pairs and genotype were treated as co-variates to take into consideration the effect *Fgf8* dosage has on embryo size.

### Apoptosis quantification

Dead or dying cells were labeled using a lysotracker red dye (L7528, thermo Fisher Scientific). Labeling of tissues were followed per manufacture guidelines. Imaging and quantification of all samples was performed together to ensure a standardized intensity of light was used for imaging between samples and that counting was standardized. Only cells localized in the mesenchyme between or next to the junction site of pp1/pc1 were quantified in the regions where pp1 first started to open until pc1 fully closed. A student’s t-test was performed to verify if the means between groups were different.

### Specimens analyzed

Sample size of evaluated specimens based on experiment are described in Table 1.

**Table 1:** Description of samples used in experiments described in this study.

| Experiment | Non-Mutant | Mutant | Age |
| --- | --- | --- | --- |
| Sox2 | 27 | 13 | E9.5-10.5 |
| Ap2-alpha | 17 | 11 | E9.5-10.5 |
| Pax1 *in-situ* | 15 | 10 | E9.5-10.5 |
| E-cadherin | 12 | 5 | E9.5-10.5 |
| F-actin | 13 | 5 | E9.5-10.5 |
| Laminin | 12 | 5 | E9.5-10.5 |
| Fgf8 in-situ | 9 | 7 | E9.5 |
| Depth Analysis | 34 | 16 | E8-10.5 |
| Shape Analysis | 34 | 16 | E8-10.5 |
| Proliferation (PH3) | 5 | 4 | E10.5 |
| Cell Death (Lysotracker) | 5 | 7 | E10.5 |

## RESULTS

### *Fgf8* reduction alters expression domains and epithelial shape

The *Fgf8* allelic series of mice consists of five genotypes that express *Fgf8* at different levels during development (Meyers *et al*., 1998). Mice heterozygous for the Neo allele (*Fgf8*^Neo/+^) express 90% of *Fgf8^+/+^* (wildtype; WT) levels, mice heterozygous for the Delta/null allele (*Fgf8^Δ/+^*) express 60% of WT levels, mice homozygous for the Neo allele (*Fgf8*^Neo/Neo^) express 35% of WT levels, and compound mutants (*Fgf8^Δ/Neo^*) express 20% of WT levels (Green *et al*., 2017). To further understand how dosage reductions impact *Fgf8* expression, we performed whole mount *in situ* hybridization in E8.5 embryos. In WT (*Fgf8^+/+^*) embryos, *Fgf8* is strongly expressed throughout the endoderm (pp1, pp2) and in the splanchnic mesoderm, whereas in the ectoderm, expression is higher on the anterior side of pc1 (Fig. 1A-C). In *Fgf8^Δ/Neo^* embryos, expression is not only reduced, but often mis-expressed. In the ectoderm, *Fgf8* extends into the proximal region where it is not normally expressed and is lost in the anterior region (red asterisk in pc1 Fig. 1D-E). *Fgf8* expression is always reduced in the mutants, however mis-expression in the cleft is variable and seems to relate to variation in severity of defects in cleft morphogenesis. We observe that the first and second cleft are often connected. *Fgf8* expression is greatly reduced in the endoderm and splanchnic mesoderm, which is typical for all *Fgf8^Δ/Neo^* embryos we examined (Fig. 1D-F).

**Figure 1:**
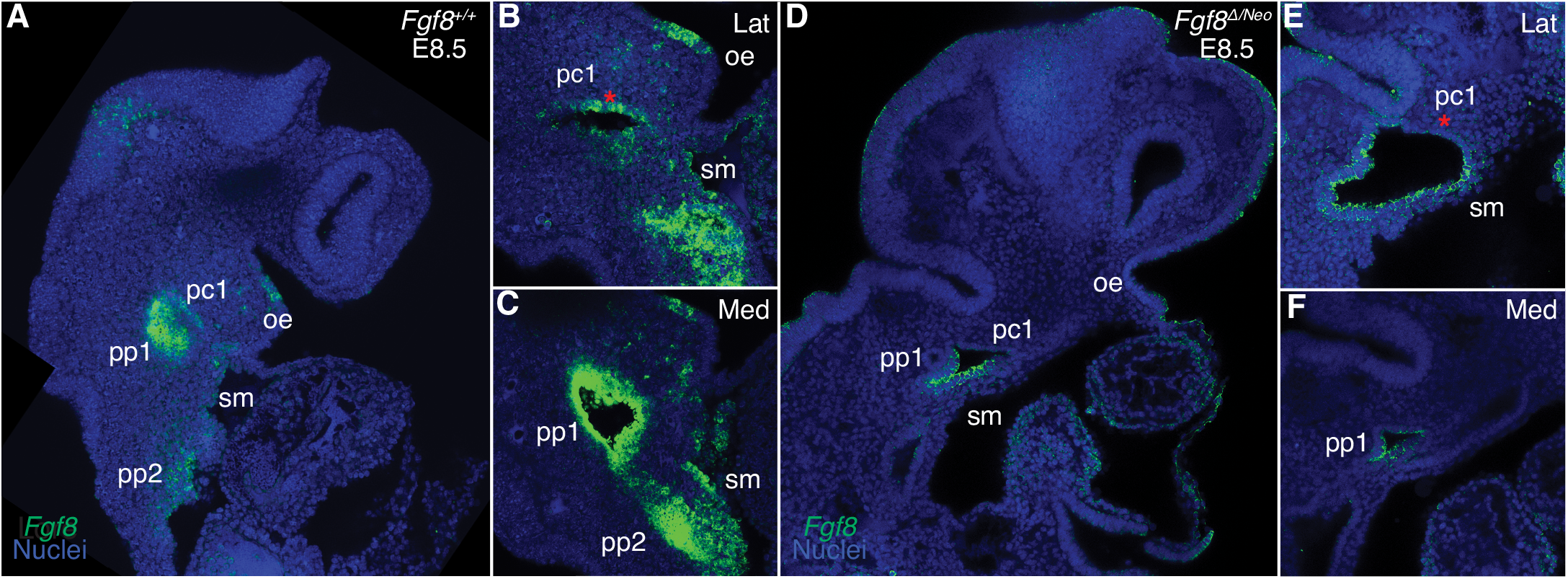
*Fgf8* is reduced and mis-expressed in *Fgf8^Δ/Neo^* epithelia. Confocal sections of whole-mount *in situ* hybridization for *Fgf8* In E8.5 embryos are shown for *Fgf8^+/+^* (WT; **A-C**) and *Fgf8^Δ/Neo^* (**D-F**) embryos. Expression is shown for an overview of the head (**A,D**) and higher magnification at a more lateral (Lat; **B, E**) and more medial (Med; **C, F**) sections. **A-C)** At E8.5, *Fgf8* is typically expressed in the oral ectoderm (oe), the ectodermal clefts (pc1), endodermal pouches (pp1, pp2), and splanchnic mesoderm (sm). **D-F)** In *Fgf8^Δ/Neo^* embryos, *Fgf8* is overall reduced in the cleft and expressed in the ventral region of pc1, but absent in the anterior region (compare red asterisk in **E** to **B**). Expression in the endoderm and splanchnic mesoderm is also greatly reduced (**F**). Note that pc1 opens widely at the lateral surface of the embryo (**E**) which often results in pc1 and pc2 being continuous in *Fgf8^Δ/Neo^* embryos.

### Out-pocketing of pp1 is largely robust to reductions in *Fgf8*

To further understand how reductions in *Fgf8* expression impact pharyngeal epithelial development, we first investigated pp1 morphogenesis. Pouch morphogenesis occurs via two separate morphogenetic processes, lateral out-pocketing and proximal-distal extension (Graham and Smith, 2001; Shone and Graham, 2014). Out-pocketing is the process where the developing pouch extends along a lateral axis from the midline to contact the overlying ectoderm (Fig. 2A). To determine how *Fgf8* dosage impacts pp1 lateral out-pocketing, we performed whole mount *in situ* hybridization for *Pax1* on E9.5 embryos and created 3D-reconstructions of pp1 from confocal images (Fig. 2B). We chose E9.5 since pp1 and pc1 are in contact and out-pocketing is nearly complete at this stage. Out-pocketing was measured as the distance from the point where pp1 first opens from the foregut medially (Fig. 2B, cool colors) to the lateral edge where it contacts pc1 (Fig. 2B, warm colors). Absolute depth is similar in all genotypes except for *Fgf8^Δ/Neo^*, which have shallower pouches (Fig. 2C; p<0.0001). However, significant lateral growth does occur in *Fgf8^Δ/Neo^* embryos, and pp1 does contact pc1.

**Figure 2:**
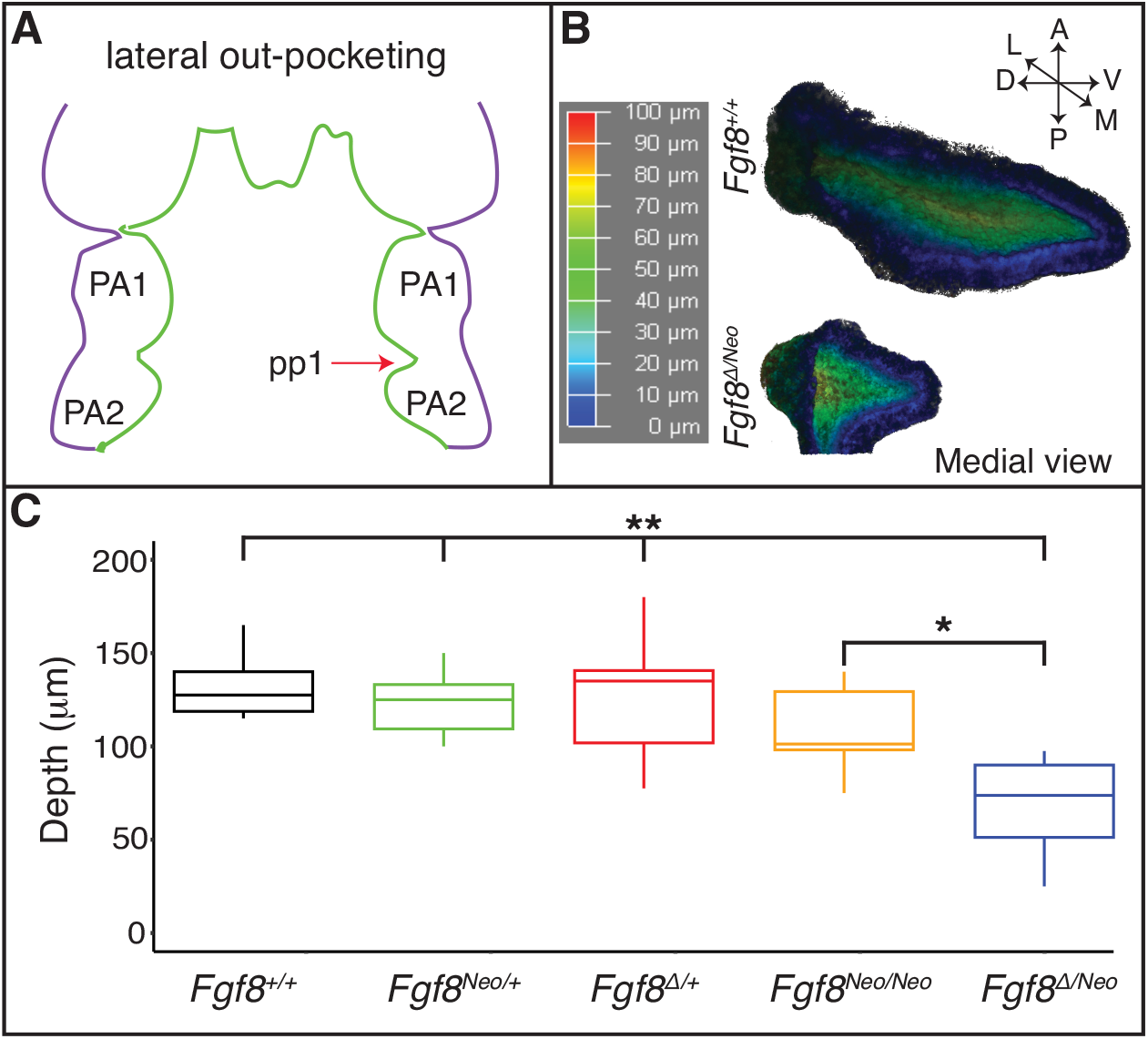
Lateral outgrowth of pp1 is robust to reductions in *Fgf8*. **A**) The first stage of pharyngeal pouch (pp1) morphogenesis is lateral outgrowth to contact the overlying ectoderm between the first (PA1) and second (PA2) pharyngeal arches. **B**) 3D reconstructions of pp1 were generated from confocal images of *Pax1* expression in the endoderm of E9.5 embryos. Cooler colors represent tissue closer to the endoderm and warmer colors represents tissue closer to the ectoderm. **C**) Only severe reductions of *Fgf8* typical of *Fgf8^Δ/Neo^* embryos result in shallower pouches (p-values <0.0001 for all genotypes compared to *Fgf8^Δ/Neo^* except *Fgf8^Neo/Neo^* where p<0.01).

### *Fgf8* is required for proximal-distal extension of pp1

To determine the effect of *Fgf8* dosage on proximal-distal extension, we quantified pp1 shape at the point where it contacts pc1 laterally for both early (12-23 somites) and late (28-31 somites) morphogenetic stages (Fig. 3). In the initial stages of out-pocketing, pp1 grows laterally as a circular tube and is circular in shape when it contacts pc1. After contact, the distal edge of pp1 extends ventrally towards the midline, resulting in a triangular shaped pouch (Fig. 3B and C upper panels). In *Fgf8^Δ/Neo^* mutants, the pouch does not extend after out-pocketing, and instead remains circular (Fig. 3B and C lower panels; see also SupFig.1). We calculated morphological disparity in pp1 shape between WT and each of the other 4 genotypes in the allelic series. Only *Fgf8^Δ/Neo^* mutants have a statistically significant difference in shape from WT at late stages (p < 0.001). We did observe, however, subtle differences in the interior shape of pp1 in *Fgf8^Neo/Neo^* mutants at E10.5, but these are not significantly different from WT (SupFig. 2). To further visualize these shape changes, we performed a Principal Component Analysis on the early- and late-stage WT and *Fgf8^Δ/Neo^* pouch shapes (Fig. 3D). Early-stage pp1 shapes occupy overlapping morphospace, which is also shared with late-stage *Fgf8^Δ/Neo^* pouches. Late-stage WT pouches differ in shape, which is observed as a shift along PC1 (representing 40% of the variation; p=0.0178). Shape changes along the PC1 axis represent alterations to pp1 reflecting compression along the anterior-posterior axis as pp1 distally extends (Fig. 3E).

**Figure 3:**
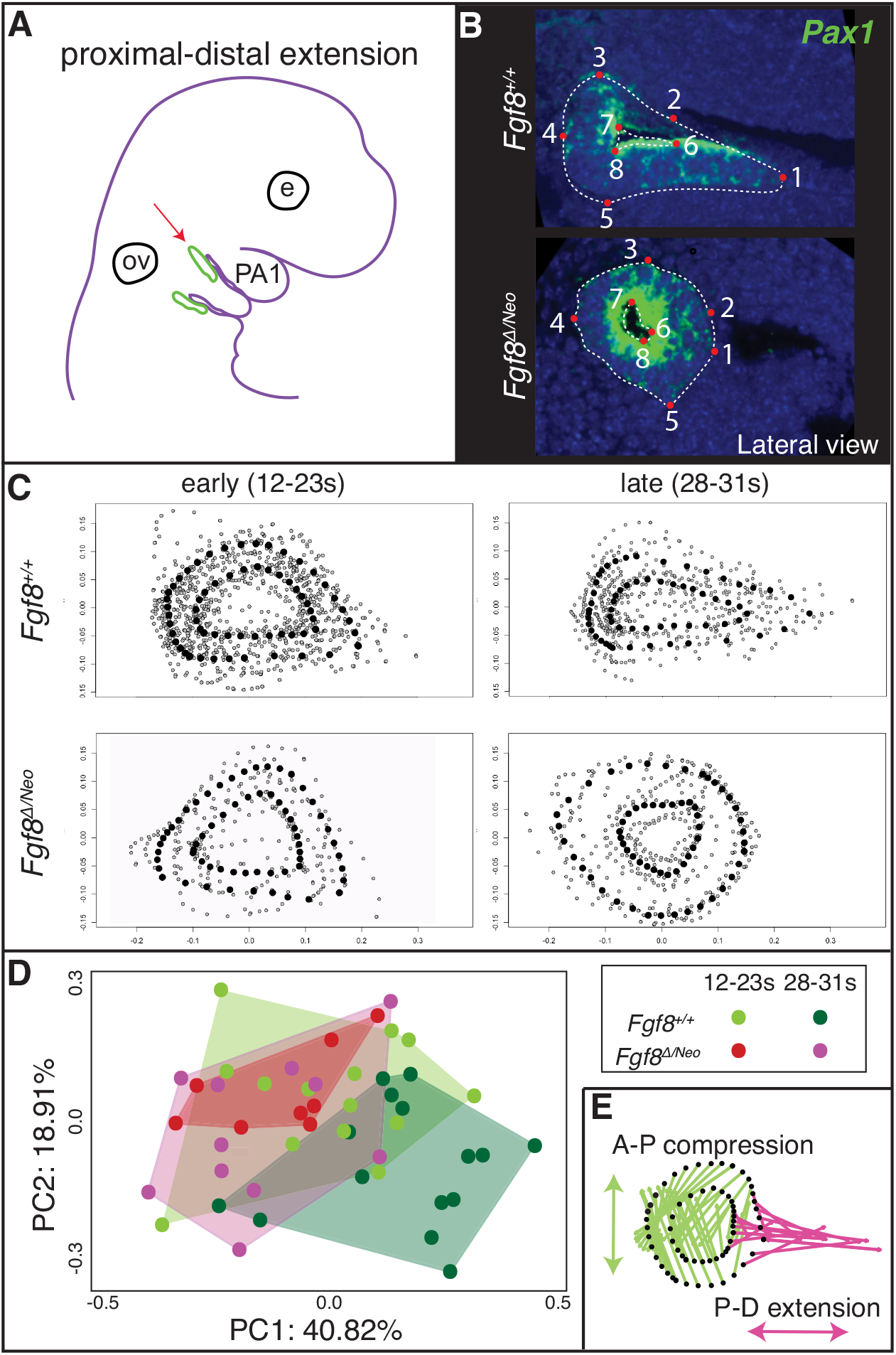
Proximal-distal extension fails in *Fgf8^Δ/Neo^* pouches. **A**) After out-pocketing, pp1 extends distally towards the jaw midline (red arrow points to pp1 in green). **B**) A landmark-based analysis was used to measure the shape and distal extension of pp1 in *Fgf8^+/+^* (WT; upper panels) and *Fgf8^Δ/Neo^* (lower panels) embryos. Eight permanent (red dots) and semi landmarks (white dashed lines) were placed on each image. **C**) Shape analysis was conducted on early (12-23 somites; left) and late stage (28-31 somites; right) embryos. **D**) A Principal Component analysis was conducted over the shape of the pouches of early (12-23 somite pairs) and late stage (28-31 somite pairs) *Fgf8^+/+^* (green) *Fgf8^Δ/Neo^* (pink) embryos. **E**) Shape change represented by PC1 is shown. The pink arrows represent distal extension. Green arrows reflect anterior-posterior compression. E, eye; ov, otic vesicle

### Staging of pharyngeal epithelial morphogenesis

To further understand how *Fgf8* reductions alter pp1 morphogenesis, we generated a standard developmental scheme for WT embryos documenting tissue interactions between pp1 and pc1 (Fig. 4). We defined 4 stages corresponding to E8.5 (pouch initiation), E9.0 (contact), E9.5 (pharyngeal plate formation), and E10.5 (pharyngeal plate modification). At E8.5 (**Pouch Initiation**), pp1 initiates out-pocketing. The foregut endoderm of pp1 extends laterally as a circular epithelial sheet towards the ectoderm (pc1). The lateral growth of pp1 occurs posterior to pc1. During the initiation stage, pp1 and pc1 are separate epithelial sheets that do not come into contact (Fig. 4A, E-G). At E9.0 (**Contact**), the lateral growth of pp1 results in contact with pc1 (Fig. 4I). At this stage, the pouch is still mostly circular in shape at the point of contact, but is more open and extended medially where it is not in contact with pc1 (Fig. 4I, J). Notably, the entire anterior side of pp1 is in contact with pc1 at its posterior-proximal edge (Fig. 4I). At E9.5 (**Pharyngeal Plate Formation**), lateral extension of pp1 continues along the proximal region of pc1 (Fig. 4L, M). At this stage, continuous contact between pp1 and pc1 occurs with the anterior side of pp1 contacting the posterior-proximal aspect of pc1. Invagination of pc1 and outgrowth of pp1 result in more overlap along the medial-lateral axis. Also at this stage, pc1 is closing at both its proximal and distal ends as the anterior and posterior ectodermal layers come in contact (Fig. 4L). This “zippering” process appears to be critical to pc1 extension. At E10.5 (**Plate Modification**), the relationship of pp1 and pc1 has changed such that the lateral aspect of pp1 is now proximal to pc1, which continues to close distally. Importantly, the proximal edge of pp1 remains in contact with pc1 (Fig. 4N-P).

**Figure 4:**
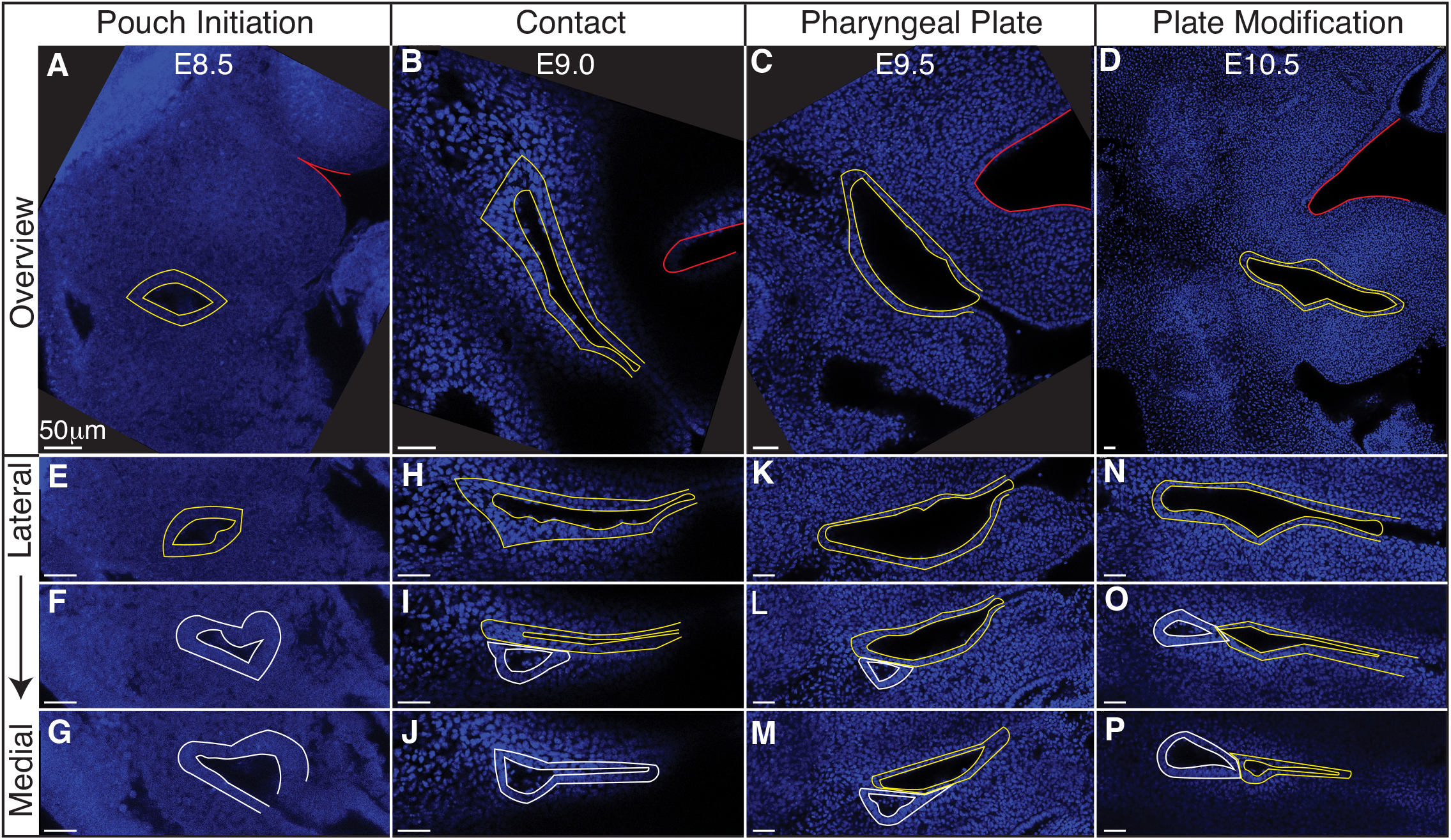
Staging of pharyngeal epithelial morphogenesis. Confocal sections from whole-mount *Fgf8* (WT) embryos are shown providing an overview of the first cleft (pc1; yellow outline) of each developmental stage at a point before the first pouch (pp1; white outline) opens in relation to the oral ectoderm (red outline) (**A-D**) and at higher magnification for three different sagittal sections from lateral to medial (**E-P**). The panels are orientated with anterior side up and ventral (distal) on the right. Each column depicts the relationship between pp1 and pc1 at 4 different morphogenetic phases: Pouch initiation, Contact, Pharyngeal Plate Formation, and Pharyngeal Plate modification. See text for more details.

### Cleft morphogenesis is abnormal in *Fgf8^Δ/Neo^* embryos

Since *Fgf8^Δ/Neo^* embryos showed significant defects in pp1 morphogenesis, we characterized pharyngeal epithelial development in *Fgf8^Δ/Neo^* embryos at the 4 stages of development described for WT (Fig. 5). Importantly, *Fgf8^Δ/Neo^* embryos exhibit highly variable defects, but some overall trends can be noted. From the earliest stage (pouch initiation) pc1 is larger and more open in *Fgf8^Δ/Neo^* embryos compared to WT (Fig. 5A,E). Contact between pp1 and pc1 often occurs earlier (already in the initiation phase) than in WT (Fig. 5F,G), which may be related to the smaller overall size of these embryos. However, this early contact does not occur in all embryos, and can even be delayed into the pharyngeal plate formation stage (Fig. 5H-J,K-M). In all cases, when contact between pp1 and pc1 occurs, it is limited temporally and spatially. Further, the position of pp1 relative to pc1 is often abnormal in the mutants. In WT, pp1 contacts pc1 at its proximal-posterior edge (Fig. 4I). However, in *Fgf8^Δ/Neo^* embryos, pp1 contact with pc1 is highly variable and can been seen anterior to pc1 or very distal along pc1 (Fig. 5L,O). Notably, during the pharyngeal plate formation stage, pp1 in *Fgf8^Δ/Neo^* embryos does not align such that its anterior side is in contact with pc1. Rather, the proximal-anterior side of pp1 is often separated from pc1 by mesenchymal cells (Fig. 5O,P; see also Fig. 6D).

**Figure 5:**
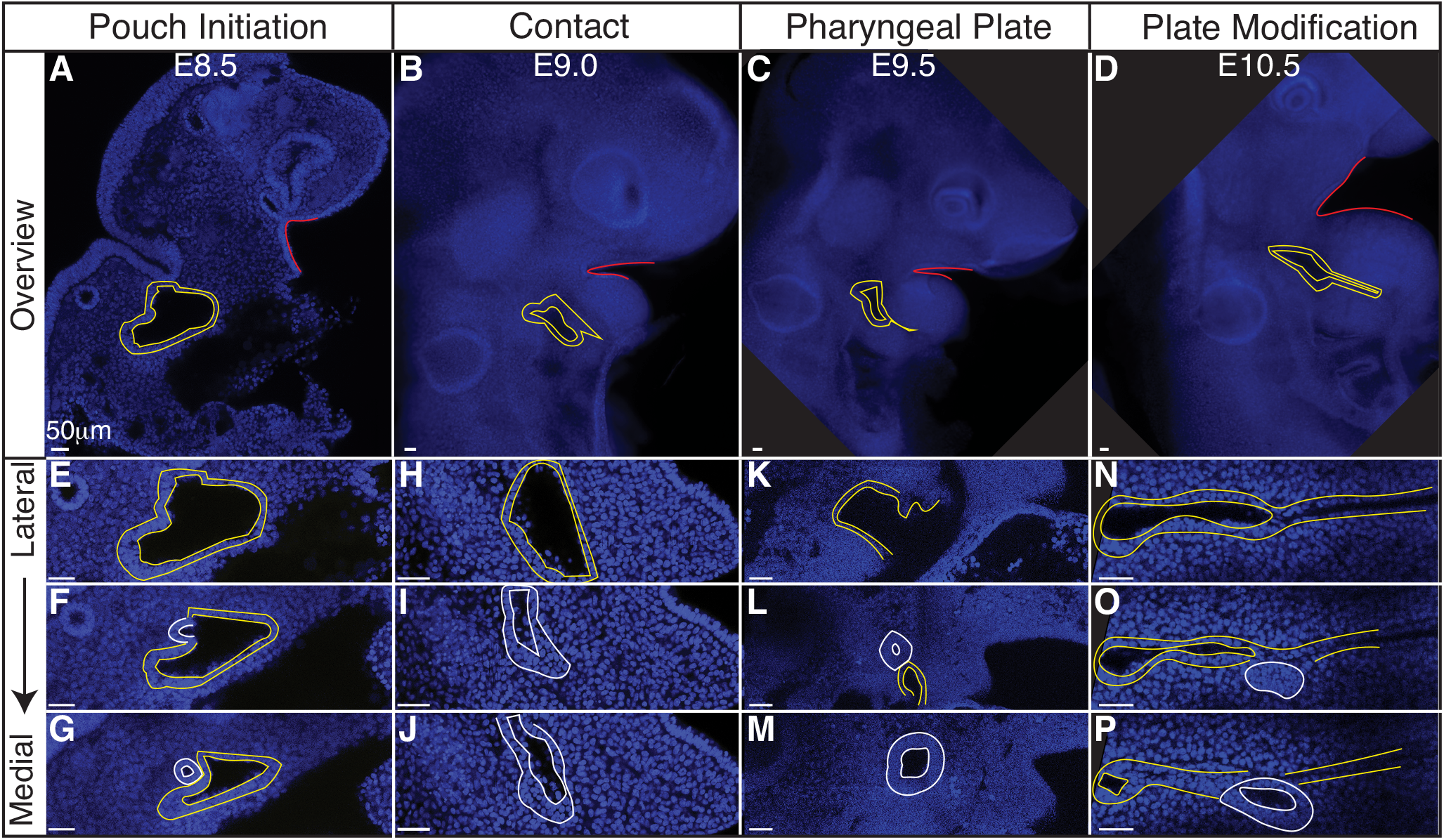
Epithelial morphogenesis is altered in *Fgf8^Δ/Neo^* embryos. Confocal sections from whole-mount *Fgf8^Δ/Neo^* embryos are shown providing an overview of the first cleft (pc1; yellow outline) of each developmental stage at a point before the first pouch (pp1; white outline) opens in relation to the oral ectoderm (red outline) (**AD**) and at higher magnification for three different sagittal sections from lateral to medial (**E-P**). The panels are orientated with anterior side up and ventral (distal) on the right. Each column depicts the relationship between pp1 and pc1 at 4 different morphogenetic phases: Pouch initiation, Contact, Pharyngeal Plate Formation, and Pharyngeal Plate modification. Notably, pc1 opens widely in early developmental stages compared to *Fgf8^+/+^* (WT) embryos. See text for more details.

**Figure 6:**
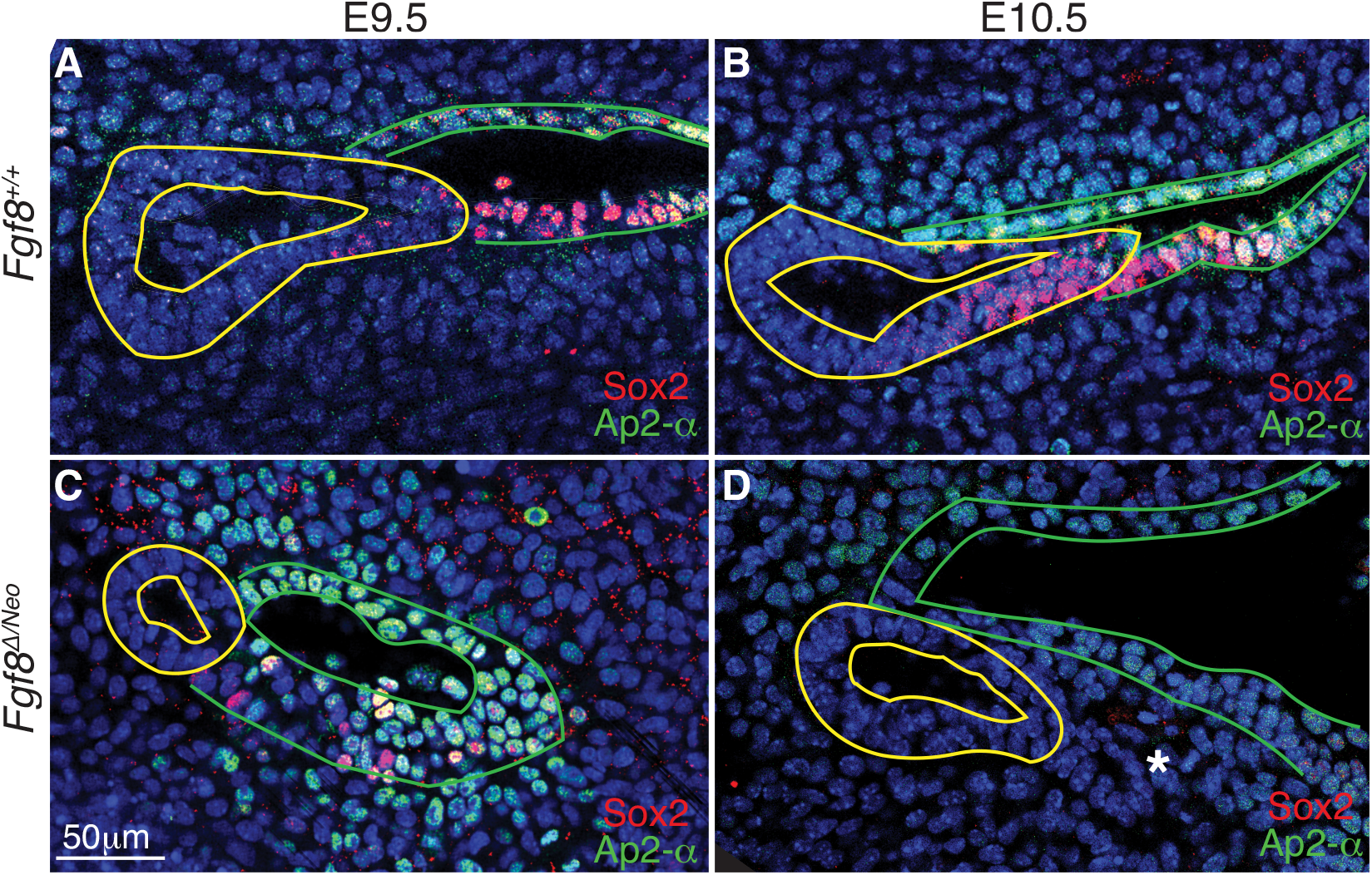
Cellular identity is altered in pp1 and pc1 of *Fgf8 ^Δ/Neo^* embryos. Confocal sections of immunostaining for Sox2 (red) and Ap2-a (green) in pp1 (yellow outlines) and pc1 (green outlines) of *Fgf8^+/+^* (WT; **A**, **B**) and *Fgf8^Δ/Neo^* (**C**, **D**) at E9.5 (**A**, **C**) and E10.5 (**B**, **D**) are shown. In *Fgf8^+/+^* (WT), all cells in pc1 are Ap2-a positive at both E9.5 and 10.5 (**A**, **B**). Additionally, Sox2 positive cells occupy the posterior-distal extending edge of pp1 and the posterior-proximal region of pc1. In *Fgf8^Δ/Neo^* embryos, pp1 lacks Sox2 expression entirely, and Ap2-a expression is much lower in pc1. In E10.5 *Fgf8^Δ/Neo^* embryos, the posterior-distal extending edge of pp1 and the posterior-proximal region of pc1 are not connected. The panels are orientated with anterior side up and ventral (distal) on the right. The white asterisk in **D** indicates a group of cells that appear to “bleb” from pp1 in this particular confocal plane.

Finally, as noted above, a normal part of pc1 morphogenesis is the process of closing, or zippering, the cleft. In WT embryos, this zippering can begin as early as E9.0 (contact) and is ongoing through E10.5 (plate modification; Fig. 4). In *Fgf8^Δ/Neo^* embryos, this zippering process fails and the anterior-posterior ectodermal layers do not come together (Fig. 5K,L,N). In many cases, pc1 remains open and contacts pc2 creating a large lateral opening (Fig. 5K). Further, because pp1 does not extend, PA1 and PA2 do not segment distally (Fig. 5D).

### Epithelial molecular identity is altered in *Fgf8^Δ/Neo^* mutants

To better characterize *Fgf8*-mediated defects in epithelial morphogenesis, we investigated cellular identity in pp1 and pc1 during pouch extension (E9.5-10.5). We performed immunostaining with Sox2, which is expressed in the foregut endoderm (Que *et al*., 2007), and AP2-alpha, which is expressed in the facial surface ectoderm (Van Otterloo *et al*., 2022). In WT, we observe that Sox2 is expressed throughout pp1 at the medial side (towards the foregut; SupFig.3A,G’), however, more laterally where pp1 and pc1 are in contact, Sox2 is only expressed in cells in the distal portion of pp1 where extension is occurring and where pp1 maintains contact with pc1 (Fig. 6A,B; SupFig.3A,E’). Notably, the proximal-posterior region of pc1 that is in contact with the distally extending endoderm is also positive for Sox2, forming a continuous region of Sox2 expression across the pp1/pc1 border (Fig. 6B). The Sox2 positive cells in the ectoderm (pc1) are also AP2-alpha positive. In contrast, Sox2 labeling is severely reduced in E9.5 *Fgf8^Δ/Neo^* pouches and completely absent in pp1 by E10.5 (Fig. 6C,D). Sox2 is also not expressed in pc1 and the region of co-expressing Sox2/AP2-alpha cells is absent (SupFig.3B,H’-J’).

### *Fgf8* mediates down-regulation of E-cadherin in pc1

Since cadherins have previously been identified as mediators of pouch morphogenesis (Quinlan *et al*., 2004), we evaluated E-cadherin localization in WT and *Fgf8^Δ/Neo^* embryos. In both WT and *Fgf8^Δ/Neo^* E9.5 embryos, E-cadherin is expressed uniformly throughout pp1 and pc1 (Fig. 7A,B). However, in WT by E10.5, E-cadherin is heterogeneous in pc1 due to localized reduction in the proximal-posterior ectoderm (Fig. 7D; SupFig.3C,G”). This area of reduced E-cadherin expression corresponds to the Sox2/AP2-alpha double-positive region described above. In contrast, at E10.5 in *Fgf8^Δ/Neo^*, E-cadherin levels remain high throughout pc1 (Fig. 7F; SupFig.3D). Notably, in WT embryos the cleft is relatively deep with a sharp slope of cells expressing E-cadherin stacked along the medio-lateral axis (Fig. 7C,E), whereas pc1 is shallow in *Fgf8^Δ/Neo^* embryos (Fig. 7G,H). Taken together, our data suggest that E-cadherin expression is mis-regulated and expanded over the surface ectoderm in *Fgf8^Δ/Neo^* embryos.

**Figure 7:**
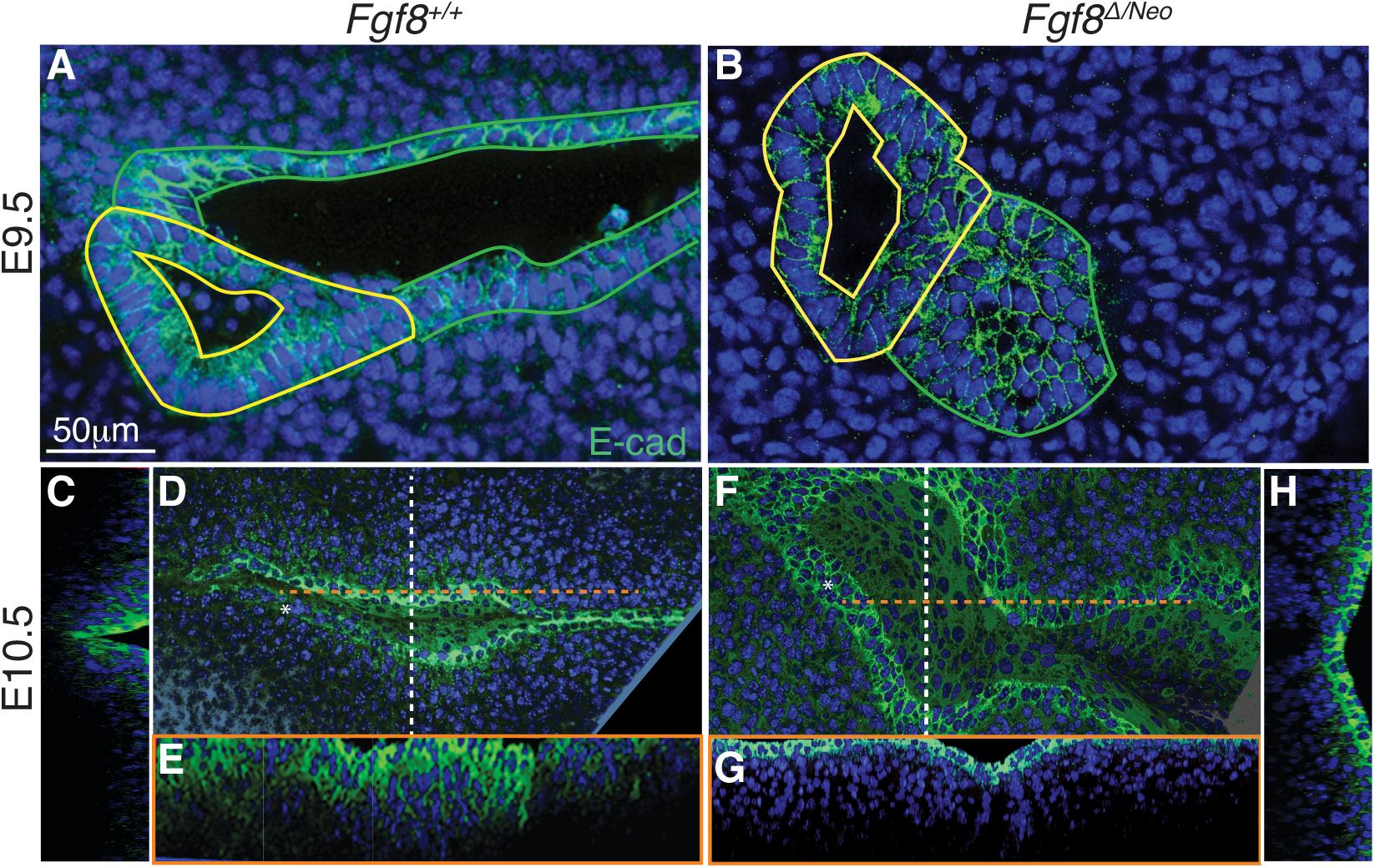
Fgf8 mediates down-regulation of E-cadherin in pc1 at the pharyngeal plate. Confocal images of E-cadherin immunostaining (green) in pp1 (yellow outlines) and pc1 (green outlines) of *Fgf8^+/+^* (WT; **A**, **C-E**) and *Fgf8*^Δ/Neo^ (**B**, **F-H**) at E9.5 (**A, B**) and E10.5 (**C-H**) are shown. At E9.5, E-cadherin is expressed throughout pp1 and pc1 in both *Fgf8^+/+^* (WT; **A**) and *Fgf8^Δ/Neo^* (**B**) embryos. **D-H**) Lateral and cross-section views of E10.5 embryos. Cross sections are color coded as dashed lines in the lateral views (**D**, **F**) as vertical (white; **C**, **H**) or horizontal (orange; **E**, **G**). The overview panels are orientated with anterior on top and distal to the right. **D**) By E10.5 in *Fgf8^+/+^* (WT), E-cadherin is down-regulated in pc1 in the proximal-posterior region (white asterisk) that connects with pp1 at the pharyngeal plate. **C**, **E**) At this stage, the cleft is closing along the proximal-distal axis, forming a deep, narrow groove. **F**) In contrast, E-cadherin remains upregulated and relatively homogenous in pc1 of E10.5 *Fgf8 ^Δ/Neo^* embryos. **G**, **H**) The cleft remains open, forming a wide, shallow groove.

### Actin cables are altered in *Fgf8^Δ/Neo^* mutants

Previous work has described that disruption to actin cables results in failure of pouch extension in chick embryos (Quinlan *et al*., 2004). To test if failure of pp1 extension in *Fgf8^ΔNeo^* embryos was associated with disruption to actin cables, we used phalloidin to label F-actin (SupFig. 4). In E9.5 WT pp1, F-actin is localized at the apical side of cells within pp1 (SupFig. 4A). However, in the anterior-distal region where it contacts pc1, Factin is more equally distributed around the cell and is also located basally (SupFig. 4B; yellow arrow). In E9.5 *Fgf8^Δ/Neo^* pp1, F-actin is highly localized on the apical side of pc1 and pp1, but no basal localization is present at the site of contact (SupFig. 4F; yellow arrow). To further assess cell polarity, we examined laminin localization. In *Fgf8^+/+^* (WT), laminin exhibits a strong basal localization in cells on the anterior side of pc1 and in cells at the proximal and anterior edges of pp1 (SupFig. 4C,D). However, in the region where pp1 and pc1 are in contact, laminin is highly expressed, but lacks basal polarization (SupFig. 4C; white arrow). In *Fgf8^Δ/Neo^* embryos, laminin is reduced, but remains polarized (SupFig. 4G,H; white arrow in G). These data suggest that loss of cell polarity is important to pp1 and pc1 interaction and that alterations to cell polarity are lost in *Fgf8^Δ/Neo^* embryos.

### *Fgf8* mediates mesenchymal cell death at the endodermal-ectodermal interface

Finally, since *Fgf8* is known to regulate cell survival and proliferation, we evaluated cell death and proliferation in E10.5 pharyngeal epithelia. We found very few pH3 positive cells in pp1 of WT or *Fgf8^Δ/Neo^* embryos (data not shown). In contrast, cell death (monitored by lysotracker staining) was prevalent (Fig. 8A-D). Notably, in WT there are numerous dying cells in the mesenchyme at the proximal region of pp1 and between pp1 and pc1 at the region of contact (Fig. 8A,C). *Fgf8^Δ/Neo^* embryos exhibit almost no cell death in this region and instead, mesenchymal cells separate pc1 and pp1 at the region of contact (Fig. 8B,D). These data suggest that mesenchymal cell death is an important component of pp1 and pc1 interaction that is lost in *Fgf8^Δ/Neo^* embryos (Fig. 8E).

**Figure 8:**
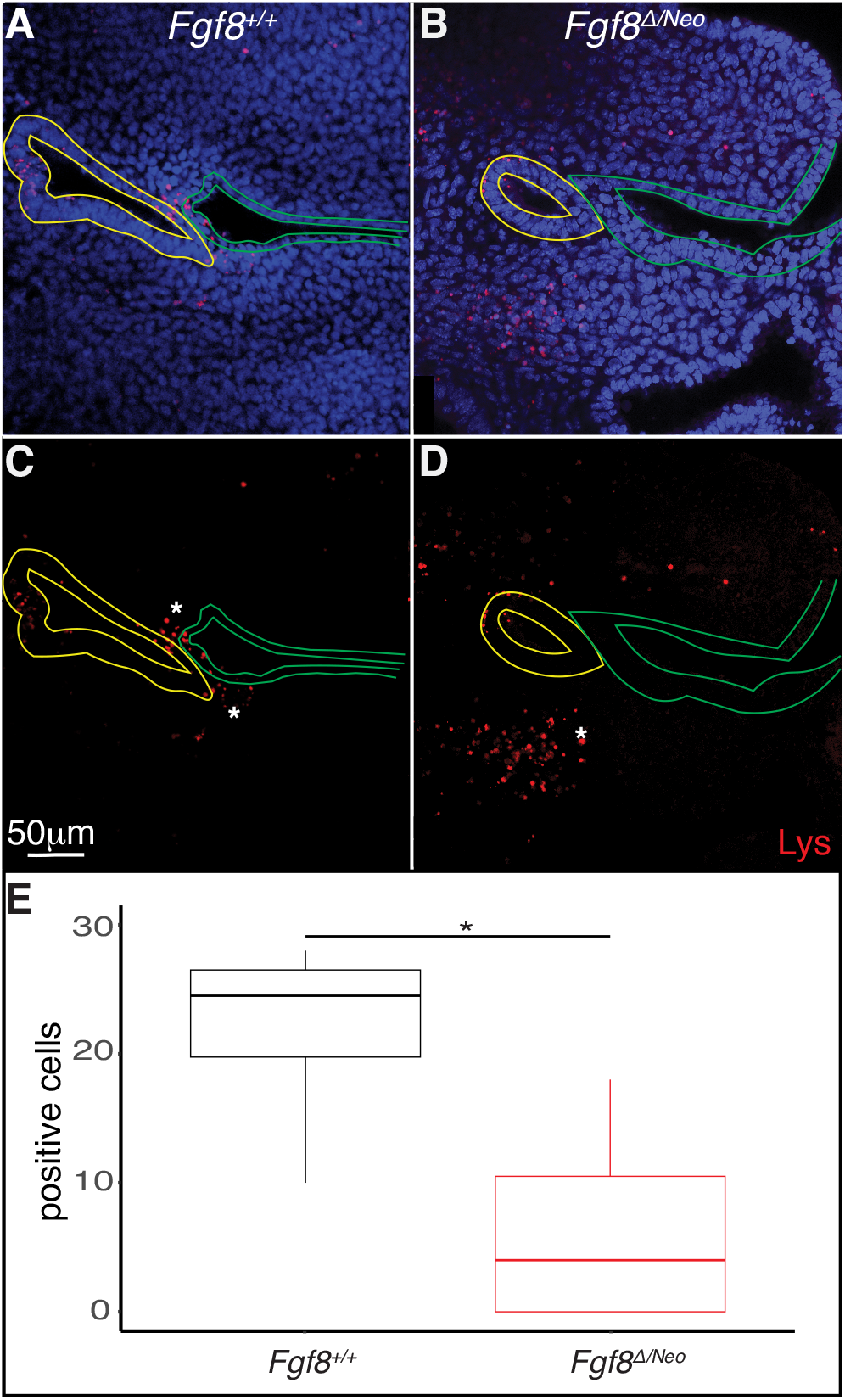
Fgf8 induces cell death in the mesenchyme surrounding the pharyngeal plate. Confocal sections of lysotracker staining (red) in pp1 (yellow outlines) and pc1 (green outlines) of *Fgf8^+/+^* (WT; **A**, **C**) and *Fgf8^Δ/Neo^* (**B**, **D**) at E10.5 are shown. Many lysotracker positive cells occupy pp1 of *Fgf8^+/+^*, but not *Fgf8^Δ/Neo^*. Additionally, many mesenchymal cells surrounding the location where pp1 and pc1 are connected in *Fgf8^+/+^*, but not *Fgf8^Δ/Neo^*, embryos. **E**) Quantification of lysotracker positive cells in the mesenchyme near the pouch-cleft connection. Student T-test; P-value = 0.0126.

## Discussion

### *Fgf8* regulates pharyngeal epithelial development

Segmentation of the pharyngeal arches is required for patterning and differentiation of the oropharyngeal skeleton. Out-pocketing of the pharyngeal endoderm between the arches is a key mediator of this process (Couly *et al*., 2002; Crump *et al*., 2004; Edlund *et al*., 2014; Graham, 2001, 2003). Although this process is essential, mechanisms regulating out-pocketing vary between the pouches and between vertebrate taxa. Notably, differences in ectodermal–endodermal interfaces between pp1 and the more posterior pouches have been described for mouse, chick, zebrafish, and shark (Shone and Graham, 2014). Here, we specifically focus on morphogenesis of pp1, in context with pc1, and the role of *Fgf8* dosage in this process in mouse embryos. We find that severe reductions of *Fgf8* levels (*Fgf8^Δ/Neo^*) disrupt both pp1 and pc1 development. Initially, pp1 does out-pocket and grow laterally to contact the ectoderm, although the total depth of pp1 is reduced compared to the other genotypes (Fig. 2). Reduction in depth may result, in part, due to reduction in size of the embryo, however, even though pp1 contacts the ectoderm, the topological relationship of pp1 to pc1 at the initial point of contact is often abnormal. Subsequently, interactions between pp1 and pc1 are altered and extension of both pp1 and pc1 is incomplete (Fig. 3). As a consequence, the first and second arches fail to separate distally.

To further understand mechanisms underlying these defects, we generated a staging series of epithelial morphogenesis in WT embryos (Fig. 4; Fig. 9). Out-pocketing of pp1 begins as a circular tube. When pp1 contacts pc1 laterally, the entire anterior side of pp1 aligns with the proximal-posterior portion of pc1. We refer to this region of contact as the pharyngeal plate (see discussion below). After contact, pp1 and pc1 extend distally together. During this process, pc1 is also elongating and “zippering” closed at both ends. Together, the connection and extension of pp1 and pc1 create a long epithelial barrier that segments PA1 and PA2 (Fig. 9). Importantly, the distal-anterior aspect of pp1 remains connected to pc1 throughout the extension process. Contact between pp1 and pc1 is mediated by reduction of E-cadherin, which occurs in a region of pc1 that is both Sox2 and AP2-alpha positive (Figs. 6,7,9). Further, a continuous sheet of endodermal and ectodermal cells share Sox2-positive identity across the region of contact between pc1 and pp1 (Fig. 6A,B).

**Figure 9:**
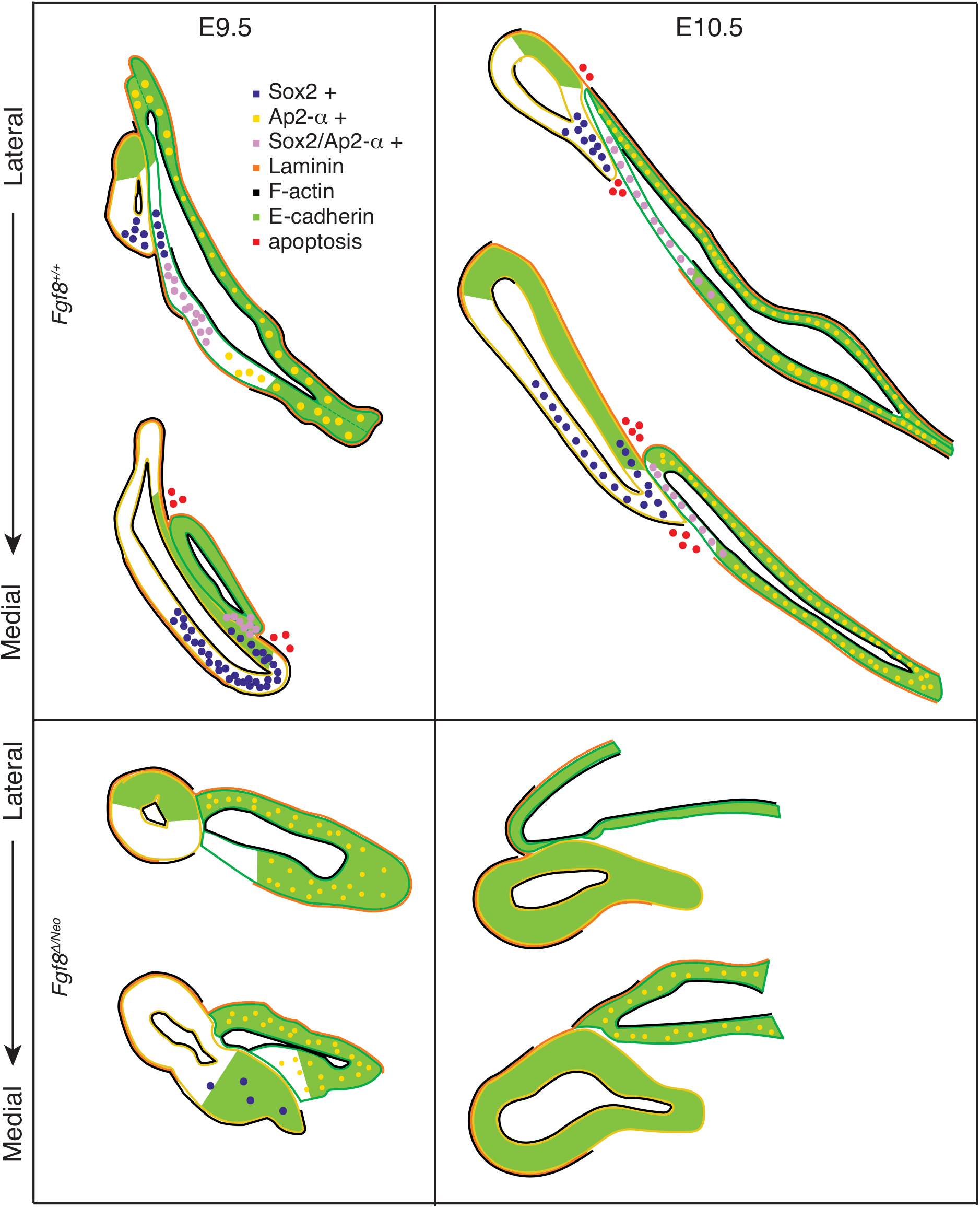
*Fgf8* regulates pharyngeal epithelial tissue interactions. Model summarizing tissue interactions between pp1 (yellow outline) and pc1 (green outline) for both *Fgf8 ^+/+^* (WT) and *Fgf8^Δ/Neo^* at E9.5 (early stage; left) and E10.5 (late stage; right) embryos. This model describes typical interactions and protein localization. For *Fgf8^Δ/Neo^*, in particular, both morphology and molecular outcomes are highly variable. See text for more details.

In *Fgf8^Δ/Neo^* embryos, these processes differ in several key aspects (Fig. 5; Fig. 9). First, the initial contact between pp1 and pc1 is highly variable and typically abnormal. Instead of forming an extended region of overlap between the epithelial layers, the proximal end of pp1 is separated from pc1 by mesenchymal cells. E-cadherin remains high in pc1 of *Fgf8^Δ/Neo^* and Sox2 expression is absent in both pp1 and pc1 (Fig. 6C,D; Fig. 7). Notably, pp1 fails to extend and instead remains circular in shape. Additionally, pc1 remains open rather than zippering closed. As neither pp1 or pc1 fully extend, PA1 and PA2 fail to segment distally.

### Tissue-specific roles for *Fgf8*

Previous work involving tissue-specific loss of *Fgf8* has shown that *Fgf8* is not required in the endoderm for pp1 morphogenesis or arch segmentation in mice (Jackson *et al*., 2014). Loss of *Fgf8* in the ectoderm using either a nestin-Cre or AP2-alpha-Cre, results in very severe phenotypes with almost complete loss of the jaw (Macatee *et al*., 2003; Trumpp *et al*., 1999). These phenotypes are more severe than those of the *Fgf8^Δ/Neo^* mutants, and in the case of *Fgf8;Nestin-Cre* embryos, were shown to be correlated with massive apoptosis in PA1 (Trumpp *et al*., 1999). Because some expression in the lateral ectoderm remains in *Fgf8;Nestin-Cre* embryos, it was interpreted that loss of *Fgf8* in the oral ectoderm drove this extreme phenotype (Trumpp *et al*., 1999). Notably, apoptosis in post-migratory NC within PA1 is not observed in *Fgf8^Δ/Neo^* embryos (Zbasnik *et al*., 2022). Mesoderm specific loss of *Fgf8* has been investigated through mesoderm (Mesp1-Cre) specific knock-out of *Tbx1* (which is upstream of *Fgf8*), as well as *Fgf8* specific deletion using Mesp1-Cre and Isl1-Cre (Park *et al*., 2006; Zhang *et al*., 2006). The focus of these mesodermal knock-out experiments was to investigate outflow tract formation, and details of first arch epithelial morphogenesis were not reported. Nonetheless, defects in pouch morphogenesis are described, with pp1 being less affected than the posterior pouches, but no information on pc1 is reported (Park *et al*., 2006; Zhang *et al*., 2006). Additionally, in zebrafish, *Fgf8* expression in the mesoderm is required for pouch out-pocketing, however, pp1 is also less affected compared to the posterior pouches and impacts on cleft morphogenesis are also not reported (Choe and Crump, 2014; Crump *et al*., 2004). In any case, our data indicate that pp1 out-pocketing is more robust to reductions in *Fgf8* than is proximal-distal extension, which fails for both pp1 and pc1 in *Fgf8^Δ/Neo^* mutants.

Notably, the mandibular phenotype in *Tbx1^-/-^* embryos is much less severe than what is observed in *Fgf8^Δ/Neo^*, rather it resembles the mildest phenotypes observed in *Fgf8^Neo/Neo^* neonates where only the coronoid process is missing (Jerome and Papaioannou, 2001; Zbasnik *et al*., 2022). Since *Fgf8* expression in pp1 and the splanchnic mesoderm, but not pc1, is regulated by *Tbx1*, this suggests that ectodermal *Fgf8* expression is a key driver behind the *Fgf8^Δ/Neo^* mandibular phenotype. Deciphering tissue-specific roles for *Fgf8*, especially separating its role in the oral vs lateral ectoderm will require more investigation. Nonetheless, our data suggest that *Fgf8* expression in the lateral surface ectoderm of pc1 is essential for arch segmentation. Importantly, in mice, arch segmentation involves extension of both pp1 and pc1. *Fgf8* is required for pc1 invagination, elongation, and zippering closed (Fig. 5; Fig. 7). *Fgf8* is also required to specify regional cell identity in both pp1 and pc1 to upregulate Sox2 and down-regulate E-cadherin, mediating their interaction and extension.

We also observe that laminin is not polarized basally in the region where pp1 and pc1 overlap. Similar observations have been made for first pouch-cleft interface in other vertebrate taxa, which were interpreted as a breakdown of the basement membrane (Shone and Graham, 2014). Despite this observation, Shone and Graham (2014) argue that no mixing of tissues occurs, with the ectoderm and endoderm remaining separate. We also find no evidence for cell mixing, but rather hypothesize this mediates interaction between the tissues for distal extension. This interaction is transient, and these layers separate by E11.5 to continue their individual differentiation (Kitazawa *et al*., 2015). Overall, our data indicate a critical role for the lateral surface ectoderm in arch segmentation that has previously been under-appreciated. A better understanding of how *Fgf8* is differentially regulated in the pharyngeal tissues, both in terms of its expression and its targets (autocrine vs paracrine) will be important to elucidate its potential impacts in disease, especially as a modifier and contributor to phenotypic variability of 22q11 deletion Syndrome.

### Boundary formation and patterning in pharyngeal development

Segmentation of the arches can also be characterized as compartmentalization, allowing separate mesenchymal cell populations to have distinct gene expression patterns and ultimately form unique skeletal elements. Compartmentalization is a critical developmental process that establishes boundaries between cell populations and also typically involves the formation of a signaling center that directs patterning after cell segregation (Dahmann and Basler, 1999; Kindberg and Bush, 2019; Pujades, 2020). The integration of pc1 and pp1 and their co-extension creates a physical boundary separating mesenchymal populations and preventing future cell mixing. We hypothesize that this contact is also critical to the formation of a signaling center organizing PA1 patterning. We have previously shown that failure to segment PA1 and PA2 in *Fgf8^Δ/Neo^* embryos is associated with alterations to the expression of patterning genes (Zbasnik *et al*., 2022). The region of contact between pp1 and pc1, referred to as the pharyngeal plate, occurs near the mid-point (intermediate region) of PA1, underlying the area that curves between the upper (maxillary) and lower (mandibular) portions of the arch.

The Hinge and Caps model of jaw development proposes that signaling interactions between the pharyngeal plate and the oral ectoderm generate a signaling center at the mid-point of PA1 (Depew and Compagnucci, 2008; Depew and Simpson, 2006). Several key lines of evidence support existence of a secondary organizer at the Hinge, or mid-point, of PA1. First, loss or gain of function experiments manipulating regulators of jaw identity (e.g., Dlx5/6, End1) result in skeletal transformations that occur as mirror-images (Depew *et al*., 2002; Gendron-Maguire *et al*., 1993; Rijli *et al*., 1993; Sato *et al*., 2008). Manipulations involving organizers typically induce mirror-image duplications because the organizer establishes a reference point for positional information (Anderson and Stern, 2016). Second, exogenous *Shh* expression near the pharyngeal plate results in duplication of juxtaposed domains of *Shh, Fgf8*, and *Bmp4* in the PA1 epithelia. As a result of this duplication of signaling interactions, the lower jaw skeleton is duplicated (Brito *et al*., 2006). Similar results have been shown for endoderm (*Shh*-expressing tissue) transplants in the pharyngeal plate region (Couly *et al*., 2002).

Our data further support the Hinge and Caps model and indicate that malformations in jaw development in *Fgf8* mutant mice result, in part, from disruptions to the jaw organizer. For proper tissue development, a signaling center must not only generate sufficient signals, but also be precisely oriented to provide accurate positional information. Our data suggest that *Fgf8^Δ/Neo^* embryos fail to form a pharyngeal plate-pp1 and pc1 have limited physical interaction and the shared molecular identity observed in WT embryos in this region is lost in mutant embryos. Therefore, appropriate signals may not be generated from these tissues. Further, severe reductions in *Fgf8* result in dysmorphic pp1 and pc1 epithelia that have altered orientations relative to the oral ectoderm. Thus, whatever signals do emenate from these epithelia will be mis-oriented.

### Evolution of the pp1 morphogenesis and origins of the jaw

The origin of the jaw is associated with an enlargement and modification of PA1 relative to the posterior arches (Kuratani, 2012). Notably, it curves near its mid-point forming an upper, maxillary and lower, mandibular portion. Similarly, pp1 is the largest of the pouches and its morphogenesis follows PA1 curvature, resulting in a pouch that is oblique, rather than parallel to, the posterior pouches. Molecular differences also exist between pp1 and the posterior pouches, which have been observed across a broad range of taxa (Choe *et al*., 2013; Crump *et al*., 2004; Okubo *et al*., 2011; Piotrowski *et al*., 2003; Quinlan *et al*., 2002). These data suggest that some aspect of pp1 development may be associated with the evolution of the jaw, which may be related to signaling interactions between pp1 and pc1 at the pharyngeal plate. In this regard, it would be important to identify conserved elements in pharyngeal plate identity among jawed vertebrates. The extent and relationship of the interaction between pp1 and pc1 requires more in-depth analysis across a broader range of taxa to determine what fundamentally characterizes this signaling center. For example, the interaction between pp1 and pc1 has been reported to be brief in chick embryos, but a detailed characterization may reveal some conserved elements (Shone and Graham, 2014).

Further, it will be important to distinguish the role of Fgfs in epithelial morphogenesis from their roles in signaling to the mesenchyme. An essential role for Fgfs in pouch formation appears to be broadly conserved among vertebrates, as FGFs drive pharyngeal pouch formation in lamprey (Jandzik *et al*., 2014). The extent to which lamprey form a pharyngeal plate is unclear, but they do form endodermal-ectodermal interfaces. The Hinge and Caps model hypothesizes that the pharyngeal plate requires interaction with the oral ectoderm. Notably, a key difference between lamprey and jawed vertebrates is the location of Fgf signaling in the oral ectoderm which is at a much greater distance from pp1 in lamprey than it is in jawed vertebrates (Shigetani *et al*., 2002). Therefore, it’s possible that the endodermal-ectodermal interface in lamprey has organizer potential but lacks appropriate interaction from the oral ectoderm. Comparisons of PA1 epithelial relationships and identity in lamprey with those from a broad range of jawed vertebrates will be required to address these possibilities.

## Acknowledgements

We would like to thank Evelyn Schwager for technical support. This work was supported by the National Institutes of Health: R03 DE028984 and R15 DE026611 to JLF.

