## Supplemental Figures for "*Fgf8* regulates first pharyngeal arch segmentation through pouch-cleft interactions"

### Zbasnik and Fish Supplemental Figure 1

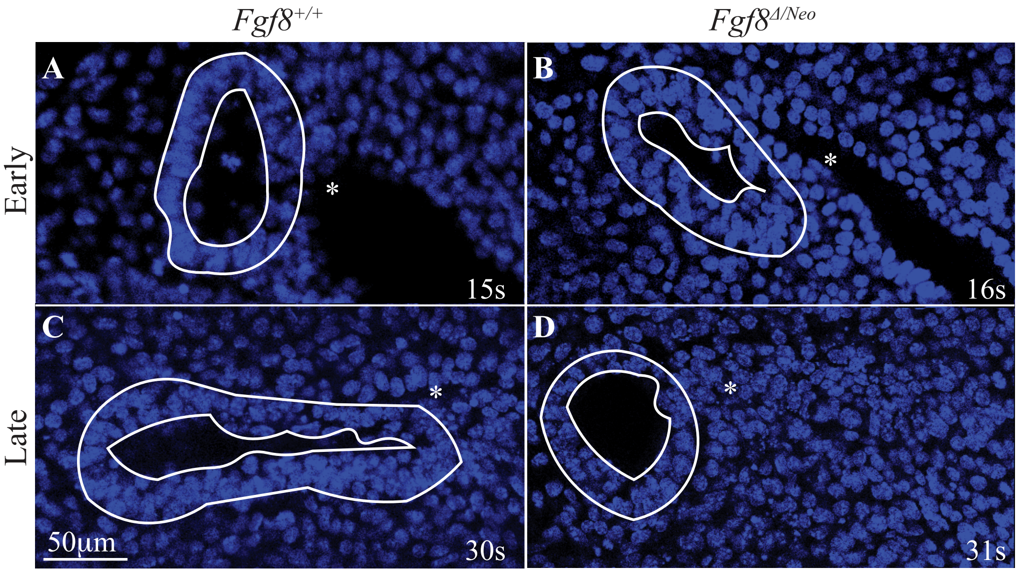


**SupFig1: *Fgf8*$^{\Delta/Neo}$ pp1 does not change shape over developmental time.** At the pouch contact stage (early), pp1 is circular in shape and is consistent across *Fgf8* genotypes as shown in **A)** *Fgf8*$^{+/+}$ (WT) and **B)** *Fgf8*$^{\Delta/Neo}$ embryos. At the pharyngeal plate stage (late), pp1 has extended distally and is triangular in shape in **C)** *Fgf8*$^{+/+}$ (WT), while in **D)** *Fgf8*$^{\Delta/Neo}$ embryos, pp1 remains circular. Asterisk indicates position of pc1.

**Zbasnik and Fish Supplemental Figure 2**

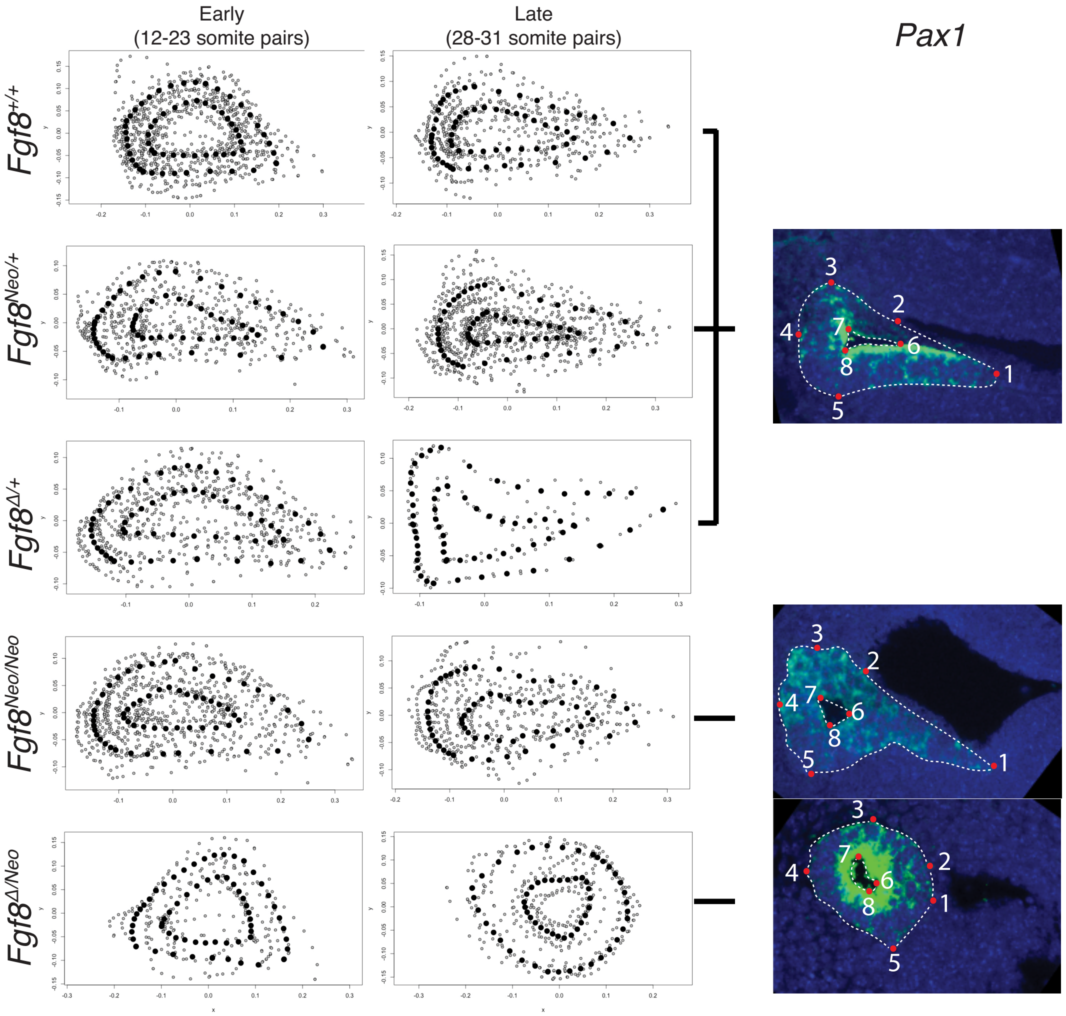


**SupFig2: Lateral shape of pp1 in *Fgf8*$^{Δ/Neo}$, but not other genotypes, differs from WT.** The average lateral shape of pp1 at early stage (12-23 somites) pouch morphogenesis is shown for all *Fgf8* genotypes (left column). The average lateral shape of pp1 at late stage (28-31 somites) pouch morphogenesis is shown for all *Fgf8* genotypes (middle column). Lateral images of *Pax1* fluorescent *in situ* hybridization and landmark placement for representative late stage embryos is shown in the right column. All panels show pp1 with anterior up and ventral/distal to the right. The red dots and overlayed numbers represent the permanent landmarks in the shape analysis where the semi landmarks are placed along the white dotted lines between each permanent landmark pair.

**Zbasnik and Fish Supplemental Figure 3**

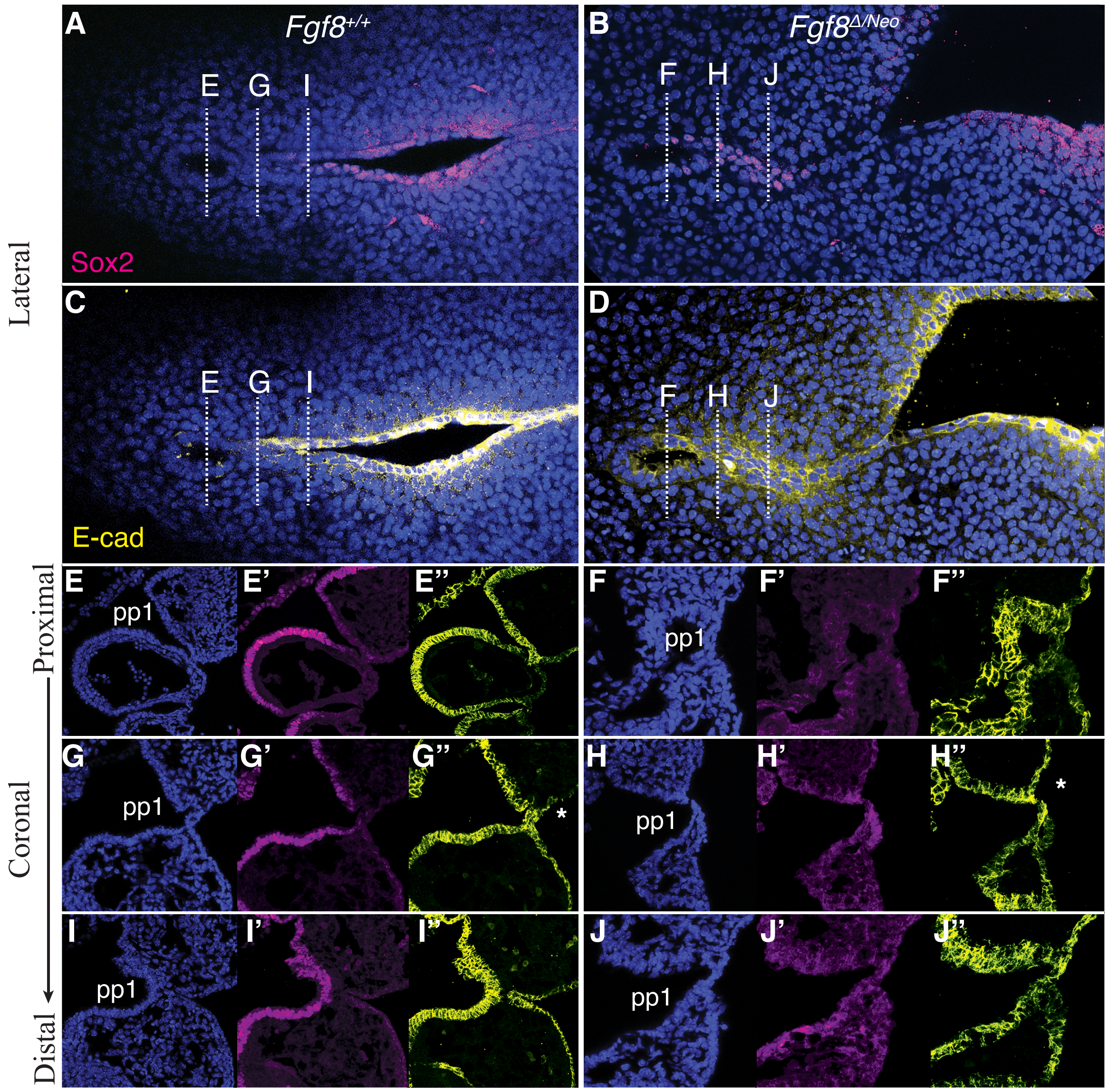


**SupFig3: *Fgf8$^{Δ/Neo}$* embryos exhibit increased E-cadherin and reduced Sox2.** Confocal sections of Sox2 (magenta) and E-cadherin (yellow) immunostaining in E10.5 embryos. A-D) Sagittal sections are shown at the plane where pp1 contacts pc1 and the pharyngeal plate in *Fgf8$^{+/+}$* (WT; A,B) and *Fgf$^{Δ/Neo}$* (C,D). These panels show pp1 with anterior up and ventral/distal to the right. E-J) Coronal sections through the first (PA1) and second (PA2) pharyngeal arches showing DNA (blue), Sox2 (magenta), and E-cadherin (yellow) immunostaining at in pp1 and pc1. These panels are cross-sections from proximal (upper) to distal (lower) cross-sections through pp1 as indicated by the white dashed lines in panels A-D. These panels are orientated with anterior up and the foregut to the left. E-G) In *Fgf8$^{+/+}$* (WT) embryos, Sox2 is low at the proximal edge of pp1 but increases distally and medially (E’,F’, G’). H-J) In *Fgf$^{Δ/Neo}$* embryos, Sox2 is low or absent throughout both epithelia and the endoderm is severely malformed (H’-J’). E-cadherin is upregulated in pc1 which is much shallower in *Fgf$^{Δ/Neo}$* embryos compared to WT embryos (H”-J”). White asterisk in G” and H” indicates region of E-cadherin down-regulation that occurs in WT, but not *Fgf$^{Δ/Neo}$* embryos.

### Zbasnik and Fish Supplemental Figure 4

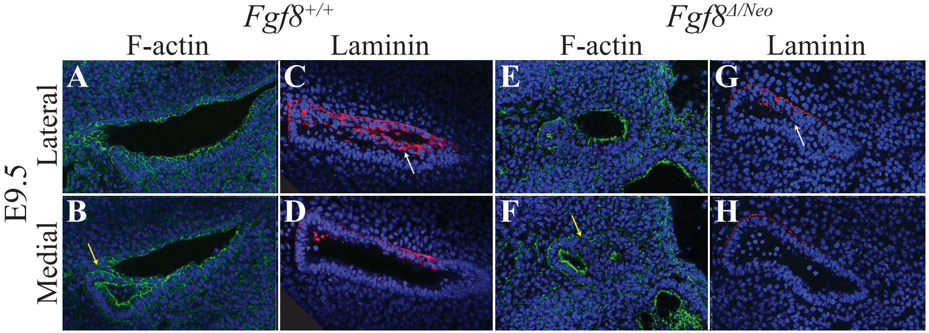


**SupFig4: F-actin and laminin are altered in *Fgf8*$^{\Delta/Neo}$ mutants.** Confocal sections of Laminin (magenta) and F-actin (green) in E9.5 *Fgf8*$^{+/+}$ (A-D) and *Fgf8*$^{\Delta/Neo}$ (E-H) embryos. All panels show pp1 with anterior up and ventral/distal to the right. In *Fgf8*$^{+/+}$ (WT), F-actin is localized apically in cells throughout pp1 and pc1, and is also located basally cells of pp1 that are in contact with pc1 (yellow arrow in B). In *Fgf8*$^{\Delta/Neo}$ embryos, F-actin is highly localized on the apical side of pc1 and pp1, but no basal localization is present at the of contact (yellow arrow in F). Instead, F-actin is located basally in cells at the proximal edge of pp1. In *Fgf8*$^{+/+}$ (WT), laminin is localized basally in cells on the anterior side of pc1 and in cells at the proximal and anterior edges of pp1 (C-D). However, in the pharyngeal plate where pp1 and pc1 are in contact, laminin is highly expressed, but lacks basal polarization (C; white arrow). In *Fgf8*$^{\Delta/Neo}$ embryos, laminin is reduced, and lacks the uniform expression in cells at the pharyngeal plate (G-H; white arrow in G).
